# S9BactDB: A database for S9 family of proteases in bacterial genomes

**DOI:** 10.1101/2025.01.01.631042

**Authors:** Soumya Nayak, Ramanathan Sowdhamini

## Abstract

S9 family proteins are serine proteases, divided into four subfamilies, involved in several functions associated with cell signalling, defence response and development. However, the annotation, characterization and statistical information are lacking. This is compounded by the huge number of bacterial genomes available. Hence, we have performed computational searches for S9 family peptidases as a step towards organising, curating the sequences and classifying into subfamilies. We have analysed annotated S9 family sequences from ∼32000 bacterial genomes/proteomes. All the curated information are presented as S9BacDB database (http://caps.ncbs.res.in/S9BactDB), provided in a user friendly way. The database provides various features such as information on the assemblies used (assembly, BioProject and BioSample details of each strain), the annotated POPs and their statistical distribution, unique domain architectures and the associated sequences and curated phylogenetic analysis. In addition, it provides the unique motifs and the associated information for Prolyl OligoPeptidases (POP) subtypes. An ML model is also integrated that can classify recognised sequences into a S9 sub family category. In conclusion, S9BactDb is a comprehensive platform that provides meticulously curated data on statistical/bioinformatics analysis of all S9 family proteins originating from fully sequenced bacterial genomes in the RefSeq database.

## Introduction

S9 family peptidase is an important family of serine proteases. Serine proteases in prokaryotic lineages are diverse enzymes with critical functions and are involved in several functions associated with cell signalling, defence response and development (1)(2). The S9 family includes important enzymes such as Prolyl Oligopeptidases (POP) and DPP IV which are implicated in various diseases. Most members of the family show restricted specificities and are active mainly on oligopeptides, consistent with the steric hindrance of the catalytic site by the beta-propeller domain at the N-terminus. For example, POP is a serine protease enzyme that specifically cleaves peptide bonds involving Proline residues (3). It is a distinct serine protease that exhibits the ability to hydrolyse internal Proline residues. According to MEROPS database, the S9 family is divided into four subcategories (4), S9A (Prolyl oligopeptidase and Oligopeptidase B), S9B (DPP IV), S9C (Acyl aminoacyl peptidase) and S9D (Glutamyl endopeptidase). Different members have different substrate specificity and thus have variety in biological significance. The active site residues are in the order Ser, Asp, His in primary sequence.

Bacterial S9 family proteins from diverse lineage have been identified and are available in the NCBI database. But the data are not complete due to incomplete annotation. Besides, they are not very convenient to use, since they are also distributed in completed and semi-completed genomes from NCBI.

Here, we report S9BactDB (http://caps.ncbs.res.in/S9BactDB), which is a comprehensive database of S9 protein family sequences. The database consists of all annotated S9 family proteins identified in all complete bacterial proteomes (available in the RefSeq database, as of February 2023). Although databases for other serine peptidases and alpha beta hydrolases exists, this is the first database specifically on S9 family sequences with additional information. Users can explore annotated POP homologues, including additional details such as strain, domain architecture, and predicted subcategories, through various browsing options and downloadable formats.

Furthermore, we offer a subcategory prediction menu that predicts the subfamily based on a given POP homologue sequence. This feature is crucial, especially considering the lack of subcategory annotations in many sequences on NCBI. We believe that this database will prove valuable in understanding the distribution and classification of diverse members within the S9 family, including insights into subfamily levels. The strain-wise statistics/information also contribute to understanding the biological significance and differences among S9 family proteins in both well-known and lesser-known species.

## Construction and content

### Datasets

The method for data acquisition is described in our previous study (5). Data from ∼32,000 complete bacterial genomes from NCBI are taken under consideration and 31,007 unique POP homologues could be identified. If we include identical homologues (100% sequence identity), the total number of annotated sequence is 52,695 (supplementary file 1). The RefSeq accession numbers, FASTA sequences, domain architecture, species/strain information, presence and absence of transmembrane helix/signal peptides, subfamily category and subfamily prediction features have been made available for the users. In addition, we have also classified the Prolyl oligopeptidases (POP, S9A) into subclusters owing to their clinical significance. Class-specific motifs were identified in our previous study suggesting that there might be differences in their substrate type and specificity. These distinct subcategories and motifs, along with the phylogenetic tree and their position in the sequence and structure, can be browsed and are downloadable.

### Implementation

Web interfaces work on client-side standard web browsers and provide easy access to various search functions. Database has been built using a standard platform based on the MERN Stack. The MERN stack is a web development framework made up of the stack of MongoDb (6), Express. js (7), React (8), and Node.js. Node.js facilitates server-side scripting with JavaScript, while NPM simplifies package management. These tools lay the foundation for building robust and scalable backends in our web application. Ubuntu Linux (version 20) is used as the operating system.

The server for prediction tool is implemented using the FLASK (9) framework with virtual environment created for Tensorflow (10) implementation of ML model in Python 3. Flask application is used to create an instance of the Flask class, enabling to define routes and configure the server.

## Utility and discussion

### Browse and Statistics menu

The database contains various pre-computed data of S9 family sequences from 32,000 complete bacterial genomes from NCBI. It provides a convenient interface for users to access the data. The homepage is shown in Fig. 1A. The desired sequence or genome assembly can be searched by users through ‘Search’ button and are browsable in different format using the browse menu (Fig. 1B).

**Fig 1:**
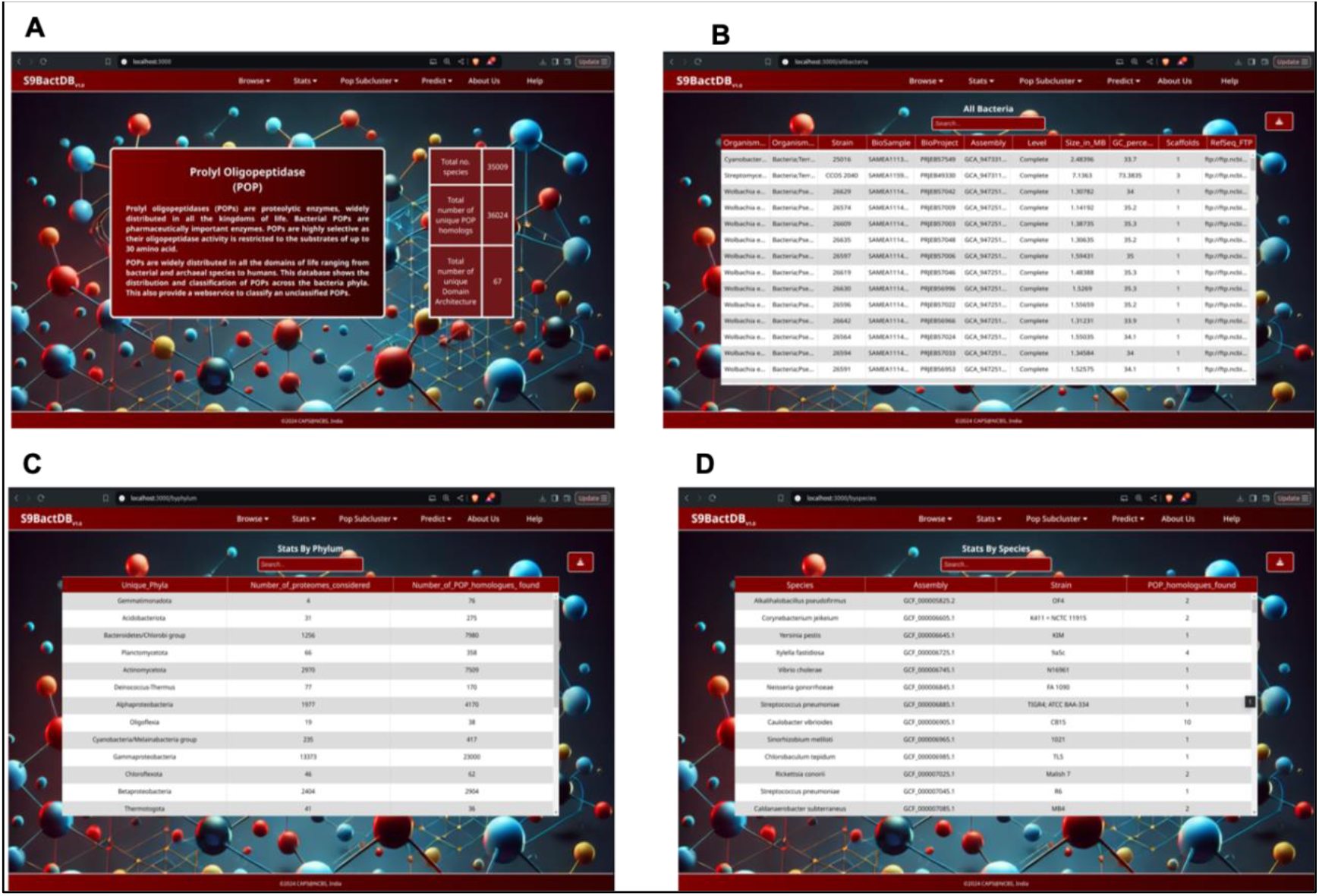
Homepage, browse and stats tab : Figure 1A represent an example representation of a. Fig. 1B shows one example from the Browse menu (Browse by all bacteria). Here the additional information of the bacterial species considered are shown. Similarly user can browse by all identified POP homologues and unique Das. The Stats menu provides statistics for each phylum, species and also the subfamily category in each species (Fig. 1C and 1D).

The browse menu consists of various tabs such as browse by genomes/assembly (with all information such as phylum, strain, RefSeq download link and BioProject) browse by accession ID of a POP homologue sequence (with all the information such as fasta sequence and the assembly it is present in) and lastly by domain architecture which presents all the possible domain architecture and the sequences in each type of domain architecture. The statistics tab presents various information on phyla and species-wise distribution (Fig 1C and D). The distribution of predicted homologues within POP subclass in each genome is also provided. In general, many genomes have only one POP representative. Few genomes have more than one POP homologues and mostly the multi-POP genomes belongs to phylum Bacteroidetes.

### POP specific Phylogenetic Tree and Cluster-specific Motifs

Within the S9 family of peptidases, Prolyl Oligopeptidases (POPs) have attracted significant research focus. These enzymes play a crucial role in neurodegenerative diseases and are also considered as potential therapeutics for celiac disease (11)(12)(13)(14). Our prior investigation delved deeply into the phylogenetic tree specific to POPs, aiming to classify it into subclusters that could give insights into substrate specificity within distinct clusters. As a result, we identified eight subclusters and associated specific motifs with each subcluster, as illustrated in Figure 2 under the tab "POP Subclusters". Users can navigate through different subclusters, and for each subcluster, sequence identity matrix can be both visualized and downloaded. This matrix offers insights into the degree of similarity or identity among sequences within a given subcluster. A heat map of the identity matrix is also provided for each subcluster, enhancing the clarity of visualization in order to facilitate user exploration. Within each subcluster, the user can view and download each of the identified motifs and their matching sequences (a motif is selected if it has a match in at least 95% of sequences). We have also identified the highly conserved positions (conserved in 90% of the sequences taking all the POPs from each clusters together). These positions are underlined in red. The matching sequences provide information about the sequence ID, exact matching sequence as well as the start and end positions. Additionally, users can access a logo featuring the associated motifs, further aiding in the interpretation of the identified patterns. This comprehensive platform offers researchers an user-friendly interface to explore and analyse the intricate details of POP subclusters, providing valuable insights into the phylogenetic relationships, sequence similarities, and conserved motifs within this enzyme family.

**Fig 2:**
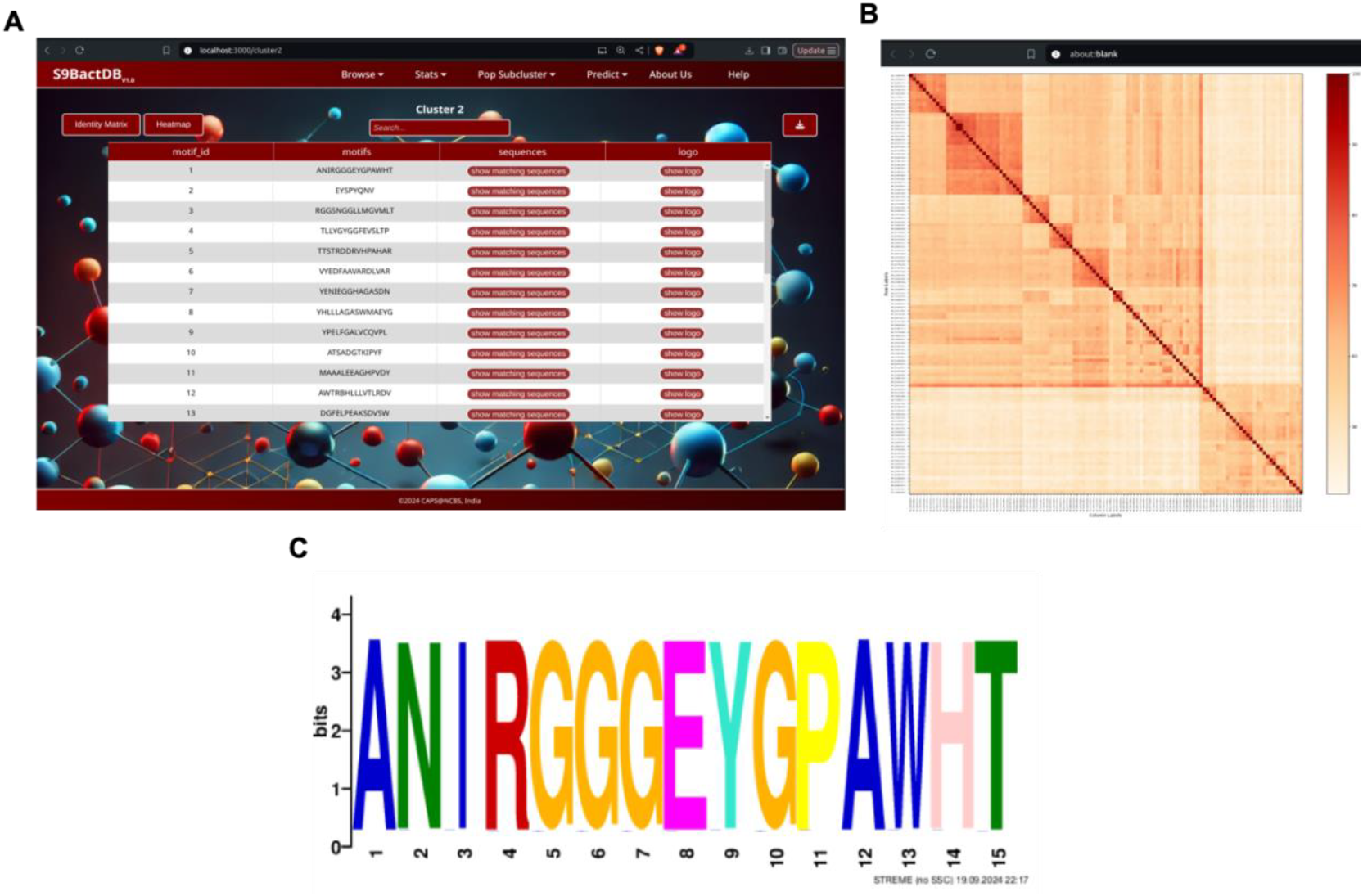
Unique motifs for POP-specific clusters. A. For each cluster the associated motifs, the matching sequences and the logo can be viewed. B.The associated heatmap of the identity matrix of a cluster sequences. C.The logo of a particular motif.

### S9 subfamily prediction console

As discussed previously, there is a lack of annotation for subfamily in the S9 family proteins in the NCBI RefSeq database. In our primary research paper (5), we undertook the subfamily classification of annotated S9 family peptidases derived from fully completed genome assemblies, totalling approximately 32,000 assemblies as of March 2023, sourced from the NCBI assembly database. To achieve this, we employed a machine learning model trained using ProteinBERT (15) and an SVM classifier. Our training dataset incorporated examples from MEROPS (4), supplemented by the limited classified subfamilies available in RefSeq. However, there are several other bacterial genomes which are nearly completed or in draft stage in RefSeq and GenBank databases (∼2,00,000). An intuitive interface has been developed to facilitate user-friendly access to our trained ML model for subcategory prediction (Fig. 3A). The backend pipeline for prediction is depicted in Fig. 3B. Only catalytic domain is considered for the training purpose, since the subfamily-specific motifs are present within this region alone according to MEROPS. At the backend, checkpoints are included to ensure that the sequence for prediction has the S9 domain. The sequence is then predicted by the already trained model for a specific subfamily (Fig. 3B).

**Fig 3:**
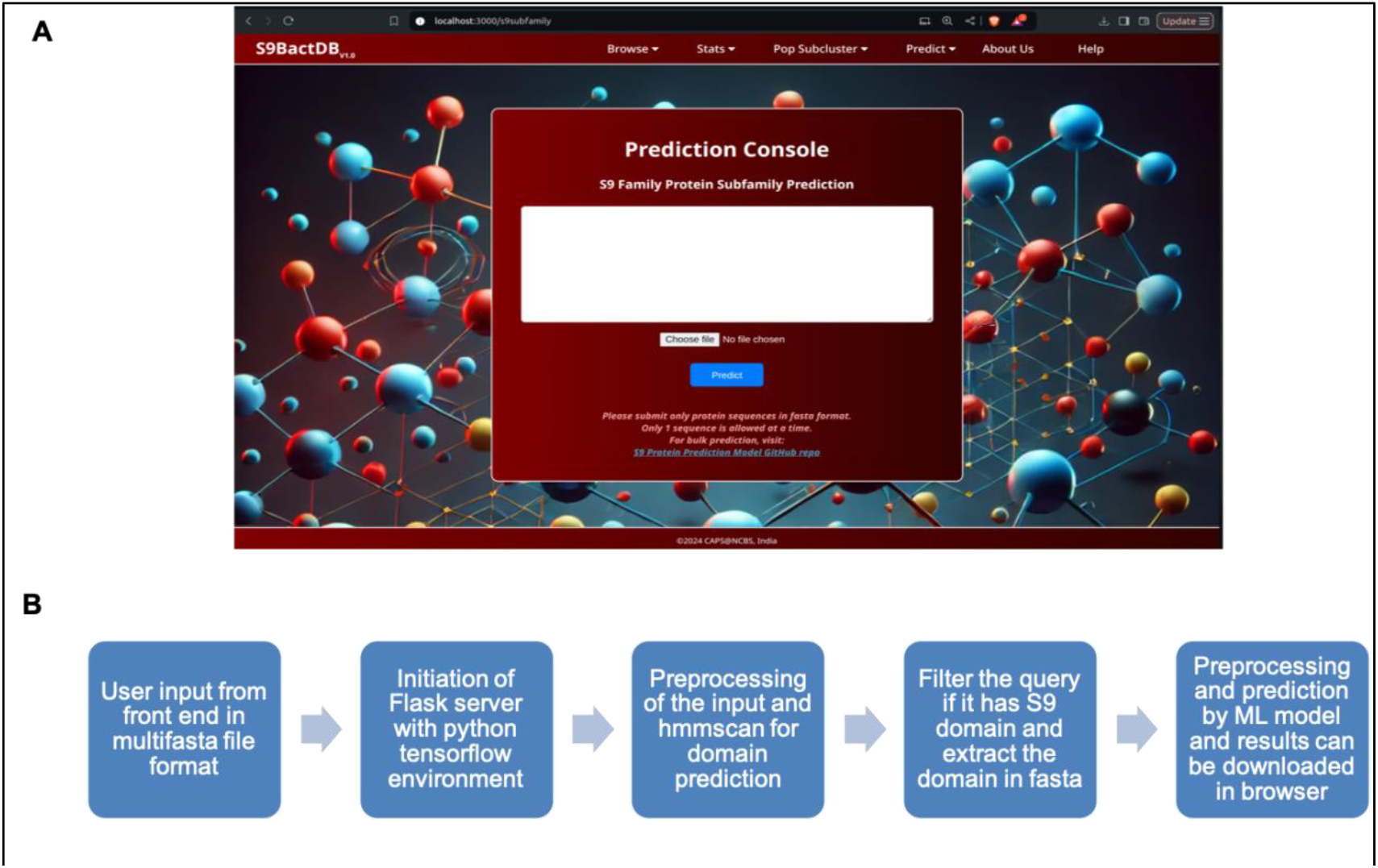
The prediction console: Figure 3A shows the prediction console. Use can paste a fasta file or browse from file system for subfamily prediction. The git hub link is provided for bulk prediction. Fig. 3B shows the pipeline used for prediction in the backend.

If the sequence is predicted/known to be a POP sequence, then there is also option for predicting the POP subcategory. For this prediction the HMM models of each subcategory is created and HMM SCAN (16)(17) is used to associate the sequence into a subcategory. E-value cut-off is used to uniquely associate a sequence (Fig 4C). This approach ensures the accuracy and reliability of our predictions for both predicting the subfamily as well as predicting the POPs and their subcategories. Due to the immense importance of S9 family in many biological applications, it is important to identify and classify a protein such that it can be assigned a biological function, this might help the biologists to address many unanswered questions. The processed and classified data from many organisms can help the users to draw homology based inferences. We believe the S9BacDB will serve as a reference for many peptidase databases aiming to organise and classify the sequences. The recorded video for navigating the website and uses of various features can be found in

**Fig 4:**
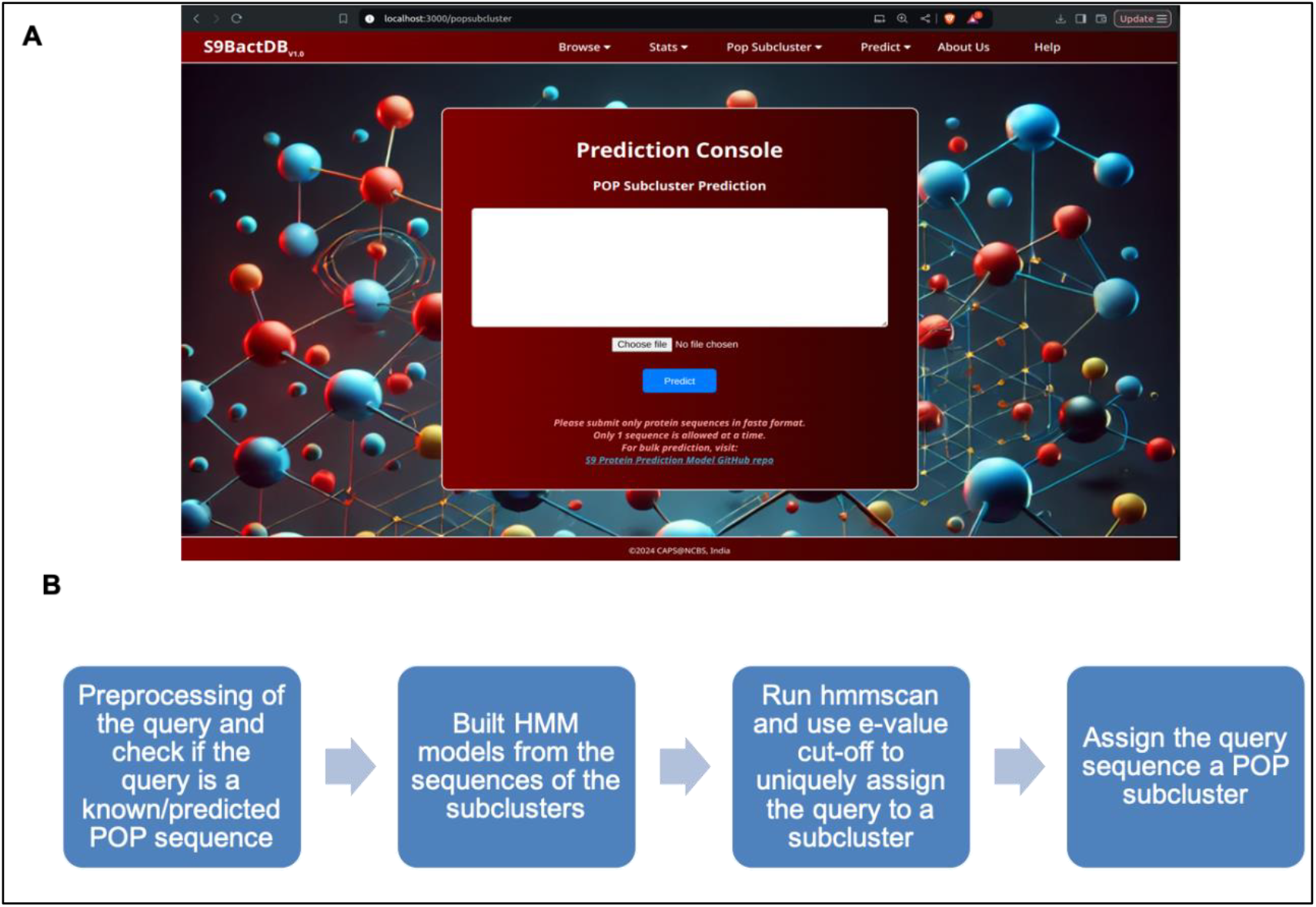
Pipeline for POP subcluster prediction. Figure 4A shows the prediction console. User can paste a fasta file or browse from file system for subfamily prediction. The git hub link is provided for bulk prediction. Fig. 3B shows the pipeline used for prediction in the backend

## Conclusion

Serine proteases in prokaryotic lineages are diverse enzymes with critical functions and are involved in several functions associated with cell signalling, defence response and development. Few subclasses, like POP, are extensively studied since they are targeted as therapeutic for celiac disease, as well as implicated in neurological disorders. The number of complete genome sequences available has exponentially increased due to the advent of next generation sequencing technologies.

S9BacDB serves as a robust platform offering curated data and statistical analyses on all S9 family proteins derived from completed genomes within the RefSeq database. Furthermore, the database provides crucial information on class-specific motifs associated with POPs. Additionally, a user-friendly webservice is incorporated to assist users in classifying POP sequences into subfamilies and assigning them to specific POP subclusters. The existing study will be further extended and involve continuous updates, incorporating cross-references to both in-house, external databases and software which can enrich the existing repository of information. It will also ensure that S9BacDB remains a comprehensive and dynamic resource for researchers delving into the intricacies of S9 family proteins and their functional implications.

## NList of abbreviations

POP: Prolyl Oligopeptidase
OPB: Oligopeptidase B
DPP IV: Dipeptidyl peptidase-4
HMM: Hidden Markov Model
ML: Machine Learning
DA: Domain Architecture

## Declarations

## Acknowledgements

The authors would like to acknowledge NCBS (TIFR), SERB, IBAB and DBT for infrastructural and other support.

## Author’s contributions

RS designed the experiments and conceived the idea. SN performed the experiments, analysed the data and wrote first draft of the manuscript and RS improved it.

## Availability of data and materials

All the data are available in supplementary material and can be browsed online.

## Consent for publication

Not applicable

## Ethics approval and consent to participate

Not applicable

